# TaxonKit: a cross-platform and efficient NCBI taxonomy toolkit

**DOI:** 10.1101/513523

**Authors:** Wei Shen, Jie Xiong

## Abstract

**Summary:** TaxonKit is a command-line toolkit for rapid manipulation of NCBI taxonomy data. It provides executable binary files for major operating systems including Windows, Linux, and Mac OS X, and can be directly used without any dependencies nor local database buiding. TaxonKit demonstrates competitive performance in execution time compared to similar tools. The efficiency, scalability, and usability of TaxonKit enable researchers to rapidly investigate taxonomy data.

**Availability:** Taxonkit is implemented in Go programming language. It is open-source and freely available for download and use from https://github.com/shenwei356/taxonkit.

## 1. Introduction

The NCBI taxonomy database is a curated classification and nomenclature for all of the organisms in the public sequence databases (NCBI, 2018). Common operations on taxonomy database include querying taxon ID (taxid) via taxon scientific names or querying lineage from taxids and so on. The online database (https://www.ncbi.nlm.nih.gov/taxonomy) provides comprehensive documents and tools for visualization and data queries.

The official Entrez programming utilities (E-utilities) (Sayers, 2010) from NCBI provides web service to query from taxonomy database, which can be used in programing ways for batch processing. Several toolkits also provide querying functions. Entrez module from BioPython (Cock, et al., 2009) wraps online Entrez services into python functions for various queries on databases including taxonomy. Theses online query tools provide access to the latest data, meanwhile, the processing speed is restricted by the Internet connection. Standalone tools parse the taxonomy dump file (ftp://ftp.ncbi.nih.gov/pub/taxonomy/) into local database to accelerate queries. The ETE Toolkit (Huerta-Cepas, et al., 2016) parses NCBI taxonomy dump files into SQLite database and provides query functions via python module. Taxadb (Gourlé, 2006) converts taxonomy data into relational databases including SQLite, MySQL and PostgreSQL. Ncbitax2lin (Xue, 2016) is a python command-line script which converts NCBI taxonomy dump file into lineages. Nevertheless, the taxonomy database updates frequently, which requires rebuilt of the local database. Besides SQL querying from relational database is not efficient especially for batch queries. Considering the small volume of taxonomy dump files and the gradually increasing read speed of storage device, directly parsing dump files is fast enough for common taxonomy data operations and free from limit of internet connection.

In this article, we describe a command-line toolkit, TaxonKit, for rapid manipulation of NCBI taxonomy data.

## 2. Method

For efficiently parsing taxonomy dump file, we choose a compiled language Go (https://golang.org/) to implement the toolkit, which supports cross-compilation for most common operating systems including Windows, Linux, and Mac OS X, and the complied single executable binary file is easy to deploy and run. Taxonkit adopts the structure of “command sub-command”, the sub-commands provide completely independent functions and all support plain or gzip-compressed inputs and outputs from either standard streams or local files. Therefore, Taxonkit can be easily combined in command-line pipes to accomplish complex operations.

At present, only nomenclature and hierarchy information in names.dmp and nodes.dmp files were used. Every sub-command of Taxonkit directly parsed and loaded necessary data into random access memory (RAM), and no daemon process is used.

## 3. Functions

### 3.1 Listing taxonomic sub-tree for given taxid

“taxonkit list” list taxids and additional information of lower taxons of given taxids, for examples, one can get the hierarchical structure of genus *Homo:*

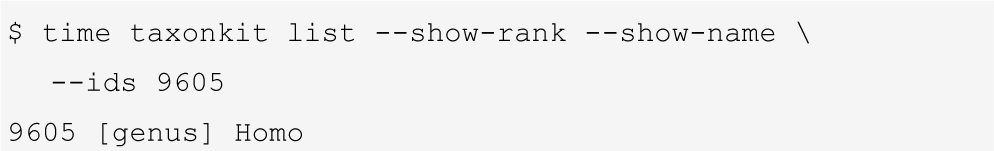

**Figure 1.**
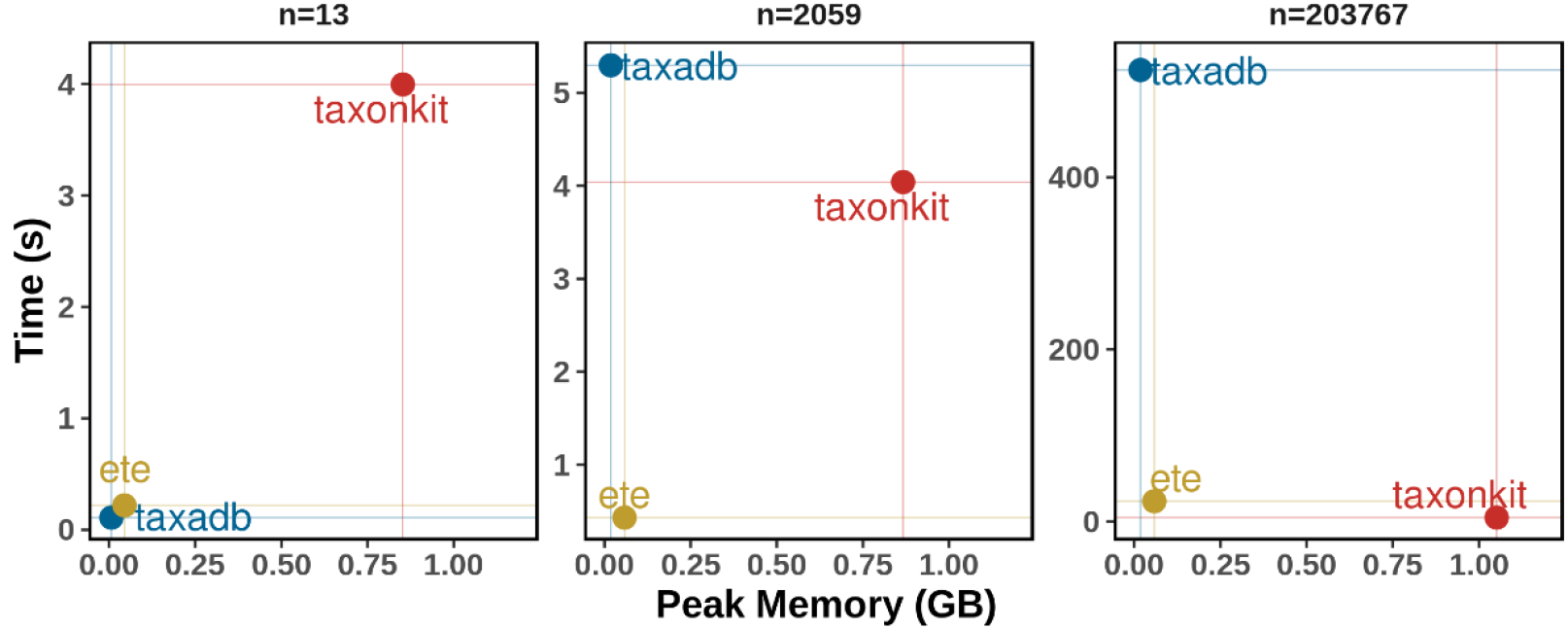
Performance comparison of querying lineage from taxids. ETE toolkit, taxadb and TaxonKit performed lineage querying for three datasets containing randomly sampled taxids from taxonomy database. All tools utilize single CPU core. Details are available on https://github.com/shenwei356/taxonkit/tree/master/bench

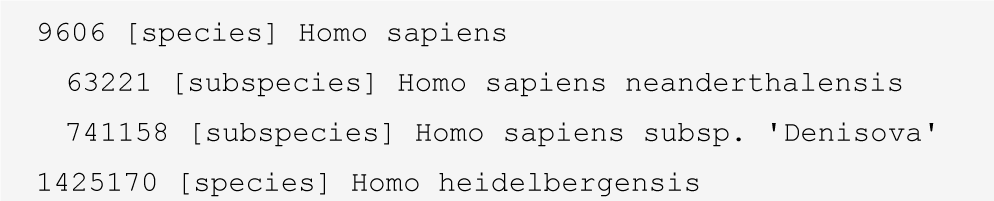

### 3.2 Querying lineage

Fetching full lineage from taxid is the most common taxonomy operation. “taxonkit lineage” command returns full lineage and corresponding taxid for given taxid. For example:

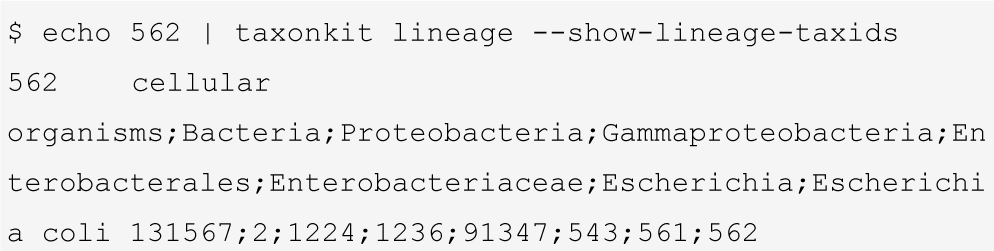

To assess the performance (execution time and peak RAM usage), Latest versions of ETE toolkit, taxadb and TaxonKit were used to query full lineage for three test datasets containing taxids randomly sampled from taxonomy database. All of these three tools utilize local taxonomy data, while Biopython was not used because it queries via internet, which is very slow for large size of queries. TaxonKit exhibited fast executing speed on different scales of data size, and the peak RAM occupation (about 1 GB) were acceptable for modern computers (Figure 1).

### 3.3 Reformatting lineage

In certain situations, user only want few ranks of the lineage instead of the verbose full lineage. “taxonkit reformat” enable users to defined ranks to retain using placeholders. For example,

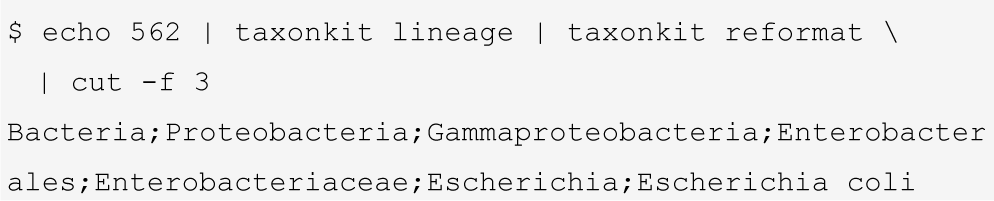

In addition, this command can estimate and fill missing rank, mainly for viruses, with original lineage information, which is useful to preprocess input data for LEfSe (Segata, et al., 2011) in visualization of phylogenetic tree. For example, lineage of taxid 11932, “Viruses; Retro-transcribing viruses; Retroviridae; unclassified Retroviridae; Intracisternal A-particles; Mouse Intracisternal A-particle” can be converted to “Viruses; unclassified Viruses phylum; unclassified Viruses class; unclassified Viruses order;Retroviridae; Intracisternal A-particles; Mouse Intracisternal A-particle”.

### 3.4 querying taxid via taxon scientific name

“Taxonkit name2taxid” is a simple command just for querying taxid via scientific names and their synonyms.

## Acknowledgements

We would like to sincerely thank all our users, especially these who reported bug and suggested new features.

## Conflicts of interest

None declared.

